# Cross-trait clustering of sub-threshold sleep genetic signals identifies EGR2 as a conserved regulator of sleep

**DOI:** 10.64898/2026.06.12.731887

**Authors:** Kyla Mace, Khanh B. Trang, Amber J. Zimmerman, Erika Almeraya del Valle, Erin N. Lottes, Rebecca E. Parker, Jakiah Choudhury, Alessandra Chesi, Struan F.A. Grant, Matthew S. Kayser

**Affiliations:** Department of Psychiatry, Perelman School of Medicine, University of Pennsylvania, Philadelphia, PA; Chronobiology and Sleep Institute, Perelman School of Medicine, University of Pennsylvania, Philadelphia, PA; Center for Spatial and Functional Genomics, Children’s Hospital of Philadelphia, Philadelphia, PA; Department of Genetics, University of Pennsylvania, Philadelphia, PA; Divisions of Human Genetics and Endocrinology and Diabetes, Children’s Hospital of Philadelphia, Philadelphia, PA; Department of Pathology and Laboratory Medicine, Perelman School of Medicine, University of Pennsylvania, Philadelphia, PA; Department of Pediatrics, Perelman School of Medicine, Philadelphia, PA; Department of Neuroscience, Perelman School of Medicine, University of Pennsylvania, Philadelphia, PA

**Author notes:** Co-first authors (Kyla Mace, Khanh B. Trang).

## Abstract

Idiopathic hypersomnia (IH) is a highly heritable sleep disorder characterized by excessive daytime sleepiness, yet few genetic pathways contributing to hypersomnolence have been identified. To expand genetic discovery beyond the limited number of genome-wide significant loci associated with sleepiness-related traits, we applied multi-trait clustering to sleep-associated genetic variation that did not reach conventional significance thresholds. Integration with our cell-type-specific variant-to-gene mapping prioritized candidate effector genes for cross-species functional screening. Among the strongest candidates was *EGR2,* which emerged as a distal effector gene at the *ADO–EGR2* locus in neurons. Neuronal knockdown of the *Drosophila EGR2* ortholog *stripe* increased sleep duration and sleep consolidation, while mutation of zebrafish *egr2* orthologs similarly increased sleep. Together, these findings demonstrate that sleep-regulatory pathways can be identified from genetic signals below conventional significance thresholds and establish EGR2 as a conserved regulator of sleep across species.

## INTRODUCTION

Idiopathic hypersomnia (IH) is a chronic neurological disorder characterized by excessive daytime sleepiness (EDS), prolonged sleep duration, and impaired arousal despite apparently sufficient sleep opportunity (*1*). In contrast to disorders of sleep fragmentation, such as insomnia or obstructive sleep apnea, IH is thought to reflect dysregulation of sleep drive itself. Patients frequently report profound sleep inertia, unrefreshing sleep, and cognitive impairment, highlighting a disruption in the processes that govern the initiation, depth, and termination of sleep. The biological mechanisms underlying IH remain poorly understood, and current treatments remain largely symptomatic rather than based on mechanism (*2*, *3*).

Genetic studies indicate that hypersomnolence is substantially heritable. Twin studies estimate the heritability of daytime sleepiness traits at approximately 40% (*4*), and a significant proportion of individuals with IH report affected first-degree relatives (*5*), However, efforts to identify genetic contributors to IH have yielded relatively few reproducible loci. The only genome-wide association study (GWAS) of IH published to date (*6*) included just 414 IH cases and did not identify any variant reaching genome-wide significance. This apparent gap between heritability and discovery reflects not only the paucity of large IH-focused cohorts but also phenotypic heterogeneity and the polygenic architecture of sleep traits. In contrast, large-scale GWAS of hypersomnia-related phenotypes, including daytime sleepiness (*7*), sleep duration (*8*), and napping (*9*), have revealed numerous associated loci. These observations suggest that genetic influences relevant to hypersomnolence are distributed across partially overlapping sleep-related traits rather than confined to a small set of disease-specific loci. Because hypersomnia-related phenotypes are difficult to capture using population-scale approaches such as single-question survey responses or actigraphy, GWAS of these traits are likely to remain underpowered. As a result, disease-relevant genetic signals may often fall below conventional genome-wide significance thresholds.

A central challenge in IH research is not simply the identification of additional loci, but the extraction of biologically meaningful signals relevant to pathological sleepiness from a broad and pleiotropic genetic landscape. Variants associated with sleep traits often influence multiple dimensions of sleep and circadian biology, complicating efforts to isolate pathways specifically related to hypersomnia. One approach to this problem is to identify groups of variants with shared patterns of association across related sleep traits, allowing biologically meaningful signatures — such as sleep fragmentation, circadian timing, or sleep propensity — to emerge from complex trait datasets (*7*, *9*).

Once such signals are identified, a second major challenge is linking the non-coding genetic variation emerging from GWAS studies to the candidate effector genes they regulate. Nearest-gene annotation is frequently misleading, as regulatory variants can act over long genomic distances through cell-type-specific chromatin interactions. Integrative variant-to-gene (V2G) mapping frameworks that incorporate chromatin accessibility and three-dimensional genome architecture provide an approach to connect genetic signals to their target effector genes in a biologically relevant cellular context. However, establishing the functional relevance of candidate genes ultimately requires experimental validation in systems where sleep can be quantitatively measured and genetically manipulated. We have previously shown that integrating variant-to-gene mapping with cross-species functional validation can translate human genetic associations into mechanistic insight. Using the few reported genome-wide significant associations for sleep propensity (*7–9*), we identified *CADM2* as a candidate effector gene (*10*), Loss of *CADM2* orthologs in *Drosophila* and zebrafish produced a hypersomnia-like phenotype, providing direct evidence that human GWAS signals can reveal conserved regulators of sleep drive across species.

While genome-wide significant associations have yielded novel hypersomnia effectors, these loci likely represent only a small subset of the broader genetic architecture underlying hypersomnia-related traits. Here, we applied Bayesian non-negative matrix factorization (bNMF) to ∼5,000 sub-threshold GWAS variants associated with related sleep traits to identify groups of variants with shared patterns of association across sleep phenotypes. Importantly, this approach extends analyses to functionally related variants not prioritized by strict GWAS thresholding, enabling the identification of a variant cluster enriched for features associated with hypersomnia including increased daytime sleepiness, prolonged self-reported sleep duration, and daytime inactivity. We then epigenetically constrained these variants using cell-specific chromatin architecture as a filter to nominate effector genes in human neurons and glia, followed by cross-species functional screening in *Drosophila* and zebrafish to identify candidate regulators of sleep. This approach converged on the transcription factor *EGR2* as a robust modulator of sleep duration and depth. Together, these findings demonstrate that coordinated patterns across polygenic sleep-trait variation can be leveraged to provide new insight into the biology of pathological sleepiness.

## RESULTS

### Multi-trait clustering identifies a sleep propensity-associated variant cluster

To implicate genetic signals relevant to hypersomnolence beyond genome-wide significant loci, we compiled sub-threshold variants (*P* ≤ 5 × 10^-^□) from large-scale GWAS of daytime napping (*9*), daytime sleepiness (*7*), and sleep duration (*8*). Following LD pruning, 4,254 independent variants were retained. For each variant, we extracted association z-scores across 18 sleep phenotypes spanning self-reported traits, accelerometer-derived measures, and obstructive sleep apnea (**Table 1, Supplementary Table 1**). We then applied Bayesian non-negative matrix factorization (bNMF) to identify groups of variants with shared patterns of association across sleep phenotypes. Across K=3-7, K=5 minimized the negative log-evidence (a Bayesian fit criterion that penalizes added clusters (*11*)) identifying it as the statistically optimal solution. (**Figure 1A,B; Supplementary Figure 1A**). However, inspection of the resulting trait profiles indicated that the K=5 solution distributed sleep efficiency and sleep duration axes across clusters that downstream functional analyses could not separate. SNP-level assignments diverged between K solutions (43 of 213 variants jointly assigned to the hypersomnia cluster under both K=4 and K=5; **Supplementary Figure 1B**), as expected when increasing K reshuffled labels. However, effector-gene annotations by V2G converged: 25 of 31 K=5 hypersomnia-related genes (80.6%) were recovered in the K=4 Cluster 2 set, which additionally nominated 68 genes that K=5 had dispersed (**Supplementary Figure 1C**). We selected K=4 as the biologically-informed solution, capturing both the K=5 gene signal and additional candidate effectors.

**Figure 1.**
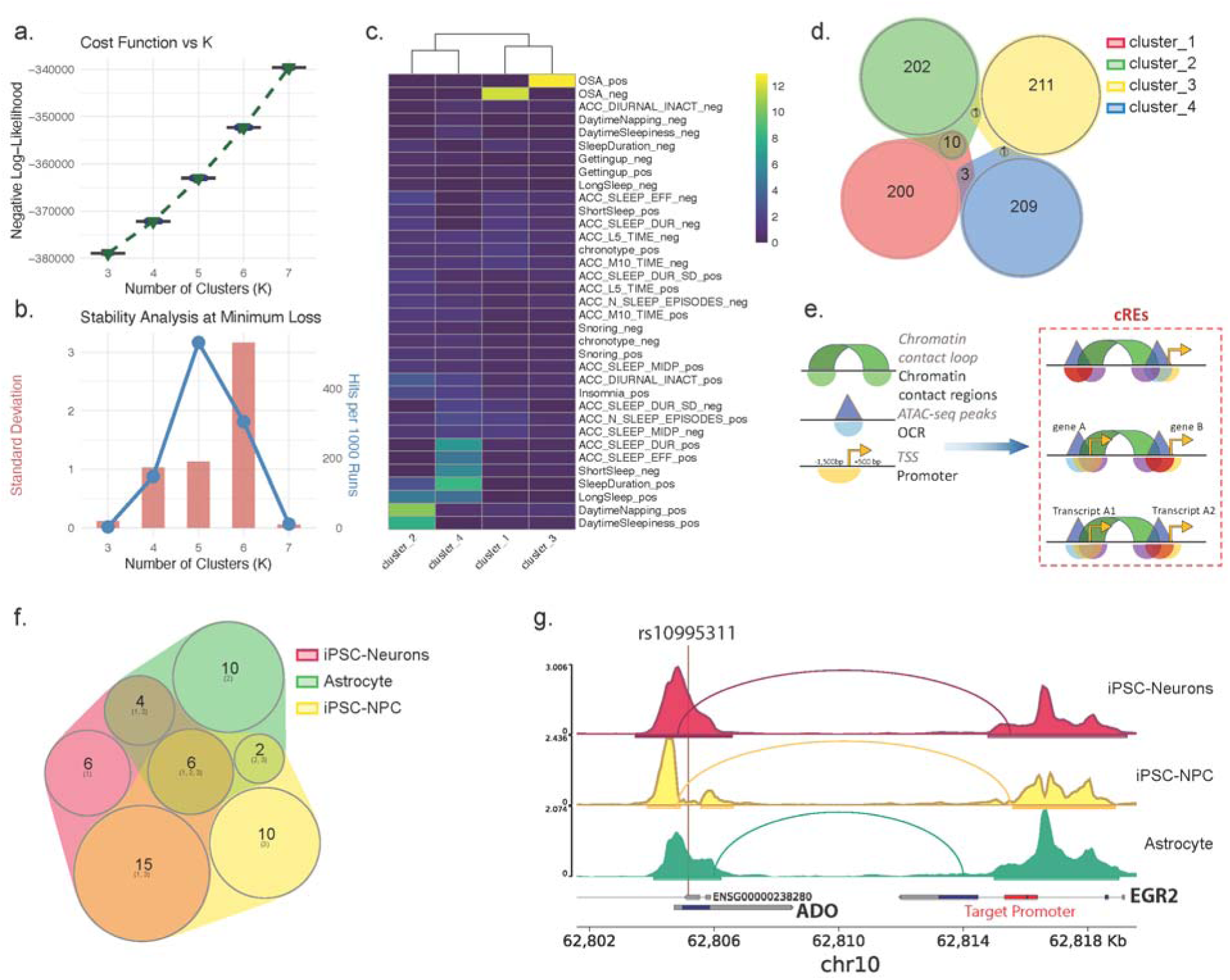
bNMF clustering of sub-threshold sleep GWAS variants to V2G mapping. (a) Model selection: negative log-Likelihood (Evidence) across K=3-7 over 1,000 runs. (b) Model selection: standard deviations of Evidence and stability across K=3-7. (c) Trait weight heatmap (H matrix): 4 clusters × 18 traits (pos/neg separated), z-scores displayed. Cluster 2 highlighted with hypersomnia-defining traits labeled. (d) Venn diagram: overlap of Cluster 2 variants between K=4 and K=5 solutions. (e) Schematic: cRE definition (open chromatin contact region vs. promoter OCR) and gene nomination logic. (f) Venn diagram: genes nominated by each cell type (iPSC-Neuron, NPC, Astrocyte). (g) Locus plots.

Inspection of sleep traits revealed that the four clusters captured distinct dimensions of sleep biology (**Figure 1C**). Cluster 1 and Cluster 3 were dominated by obstructive sleep apnea signals, while Cluster 4 was associated with objectively measured long sleep duration and increased sleep episodes without accompanying daytime sleepiness, consistent with a constitutive long-sleeper phenotype. Cluster 2 showed positive associations with daytime sleepiness, daytime napping, prolonged self-reported sleep duration, and accelerometer-measured daytime inactivity, identifying a group of variants enriched for features associated with increased sleep propensity.

Using a 95th percentile cluster-weight threshold, we identified 213 variants in Cluster 2 as candidate hypersomnolence-associated loci (**Figure 1D; Supplementary Table 2**). Notably, all six previously described “sleep propensity” loci (*9*) were assigned to Cluster 2 by the 95th-percentile weight threshold, supporting the validity of this clustering approach.

### Cell-type-specific V2G mapping identifies candidate hypersomnolence genes

To link Cluster 2 variants to candidate effector genes, and to apply a meaningful biological filter to the subthreshold nature of these signals, we performed cell-type-specific variant-to-gene (V2G) mapping (*12*) using cis-regulatory elements (cREs) annotations defined by open chromatin regions (OCRs) and three-dimensional chromatin interactions in human neural cell types (**Figure 1E**). We leveraged our chromatin accessibility maps from induced pluripotent stem cell (iPSC)-derived cortical neurons (*13*), neural progenitor cells (NPCs) (*13*), and astrocytes (*14*) and integrated these datasets with promoter Capture-C maps to connect non-coding variants to putative target genes. A gene was nominated when a chromatin loop connects a variant-containing cRE to a target gene’s promoter OCR. Because regulatory variants frequently act over long genomic distances rather than on the nearest annotated gene (*15*), this epigenetically-informed approach enabled assignment of Cluster 2 variants to candidate genes within cell-type-specific regulatory networks.

Across these datasets, V2G mapping nominated 93 targets from the 213 Cluster 2 variants, including 53 protein-coding genes (**Figure 1F**; **Table 1**). Approximately half of the nominated genes were found in more than one cell type (example in **Figure 1G**), while others showed cell-type specificity. The astrocyte chromatin maps identified 10 additional candidate genes not captured by neuronal datasets alone. Together, these analyses generated a prioritized set of candidate hypersomnolence genes for subsequent cross-species functional validation.

### Functional screening in Drosophila identifies stripe/EGR2 as a conserved sleep regulator

To functionally evaluate V2G-nominated candidate genes, we performed a cross-species RNAi screen in *Drosophila melanogaster* using neuronal (*elav*-GAL4) and glial (*repo*-GAL4) drivers (**Figure 2A,B**). We used reagents from multiple RNAi collections (*16*, *17*) to knock down fly orthologs of candidate hypersomnolence genes and quantified sleep using the *Drosophila* Activity Monitoring (DAM) System (Trikinetics). Because the DAM system produces multiple correlated measurements of sleep duration, sleep architecture, and waking activity, we used principal component analysis (PCA) followed by *k*-means clustering to identify genotypes with shared behavioral phenotypes across the screen. The clusters captured distinct genotype-associated phenotypes (**Figure 2C,D**), including altered waking activity with unaffected sleep (neuronal Cluster 2), sleep fragmentation (neuronal Cluster 4, glial Cluster 2 and 4), broad locomotor deficits (glial Cluster 3), and minimal behavioral effects (neuronal Cluster 3). Together, these patterns highlight the phenotypic diversity of effector genes linked to the human sleep propensity variant set. However, in both neuronal and glial datasets, a single cluster (Cluster 1) was characterized by increased sleep duration, longer sleep bouts, and minimal reductions in waking activity, consistent with an elevated sleep propensity phenotype (**Figure 2C-F**). The neuronal screen produced stronger sleep-promoting phenotypes than the glial screen, with more genotypes at the extremes of the PCA space (**Figure 2E,F**).

**Figure 2.**
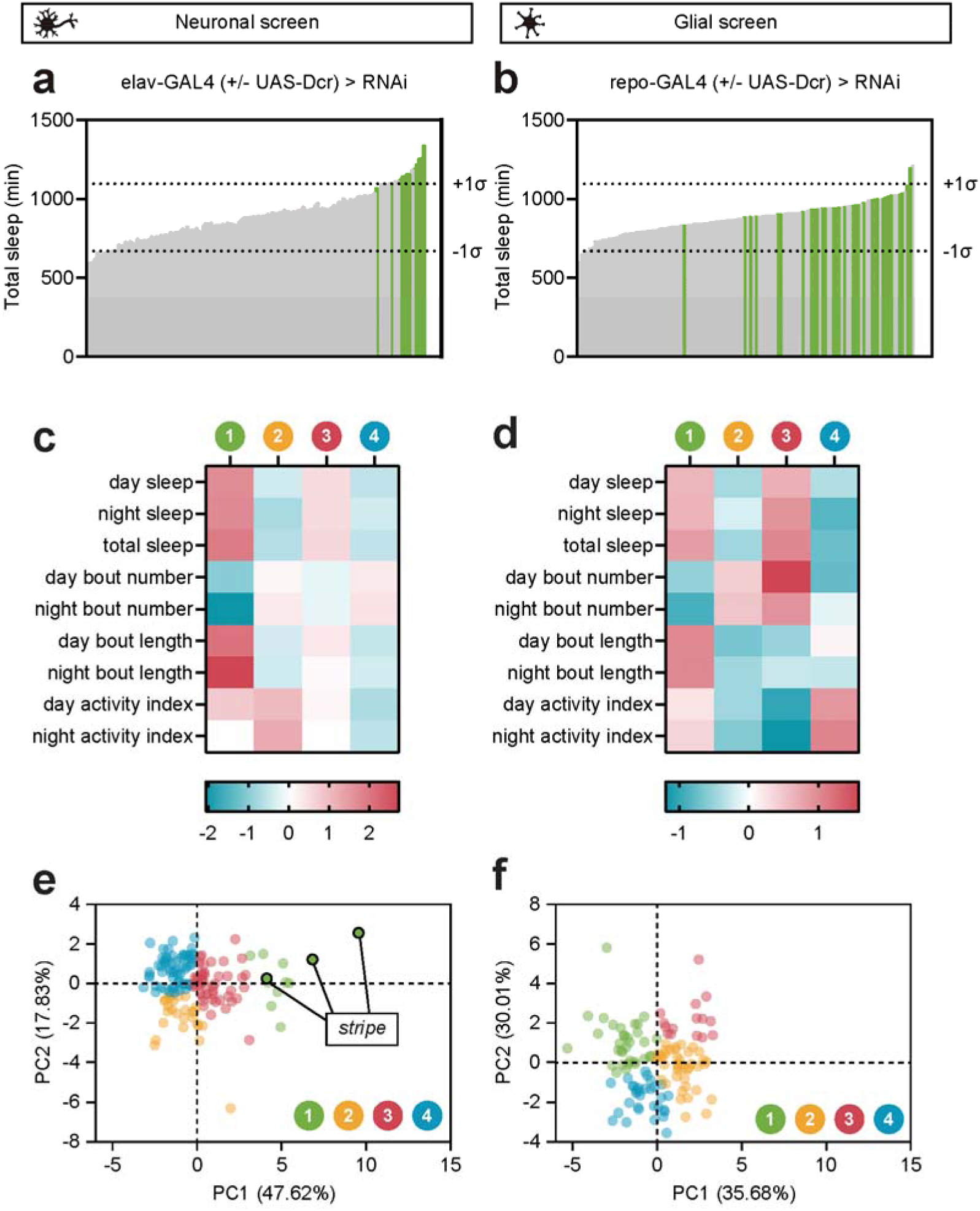
Reverse-genetic RNAi screen of hypersomnolence-associated genes reveals clusters of genotypes with hypersomnia-like phenotypes. (a, b) Total sleep across 24 h of flies expressing RNAi driven by pan-neuronal elav-GAL4 (a) or pan-glial repo-GAL4 (b). UAS-Dcr2 was co-expressed with RNAi of the VALIUM1, VALIUM10, or VDRC vectors to improve RNAi efficacy. Bars indicate the mean sleep measurements across 8-16 flies, except for 3 elav-GAL4 lines which have 5-7 animals and 2 repo-GAL4 lines which have 4 and 7 animals. Green bars represent genotypes captured in the hypersomnia-like cluster, described in (c-f). (c,d) Heat map representation of sleep phenotype clusters identified from PCA/k-means clustering. Cluster 1 defines a group of hypersomnia-like genotypes, exhibiting longer sleep duration, fewer sleep bouts, longer sleep bouts, and minimal effects on activity index. Heatmap displays the mean value of the sleep measure within each cluster, and measures were standardized such that the overall mean for each measure is 0 and the standard deviation is 1. (e, f) PCA plot of neuronal screen, color-coded by clusters identified in (c, d). Three long-sleeping genotypes expressing RNAi against *stripe* are highlighted.

Among the strongest hits were three independent RNAi lines targeting a transcription factor *stripe* (*sr*), the fly ortholog of EGR2, all of which produced robust increases in sleep. Replication in a higher resolution sleep monitoring system confirmed that neuronal *stripe* knockdown increased daytime and nighttime sleep duration and consolidation without reductions in waking activity (**Figure 3; Supplementary Figure 2**). Because *EGR2* emerged as one of the most compelling candidates from the functional screen, we next returned to the human genetic data to further investigate the implicated locus.

**Figure 3.**
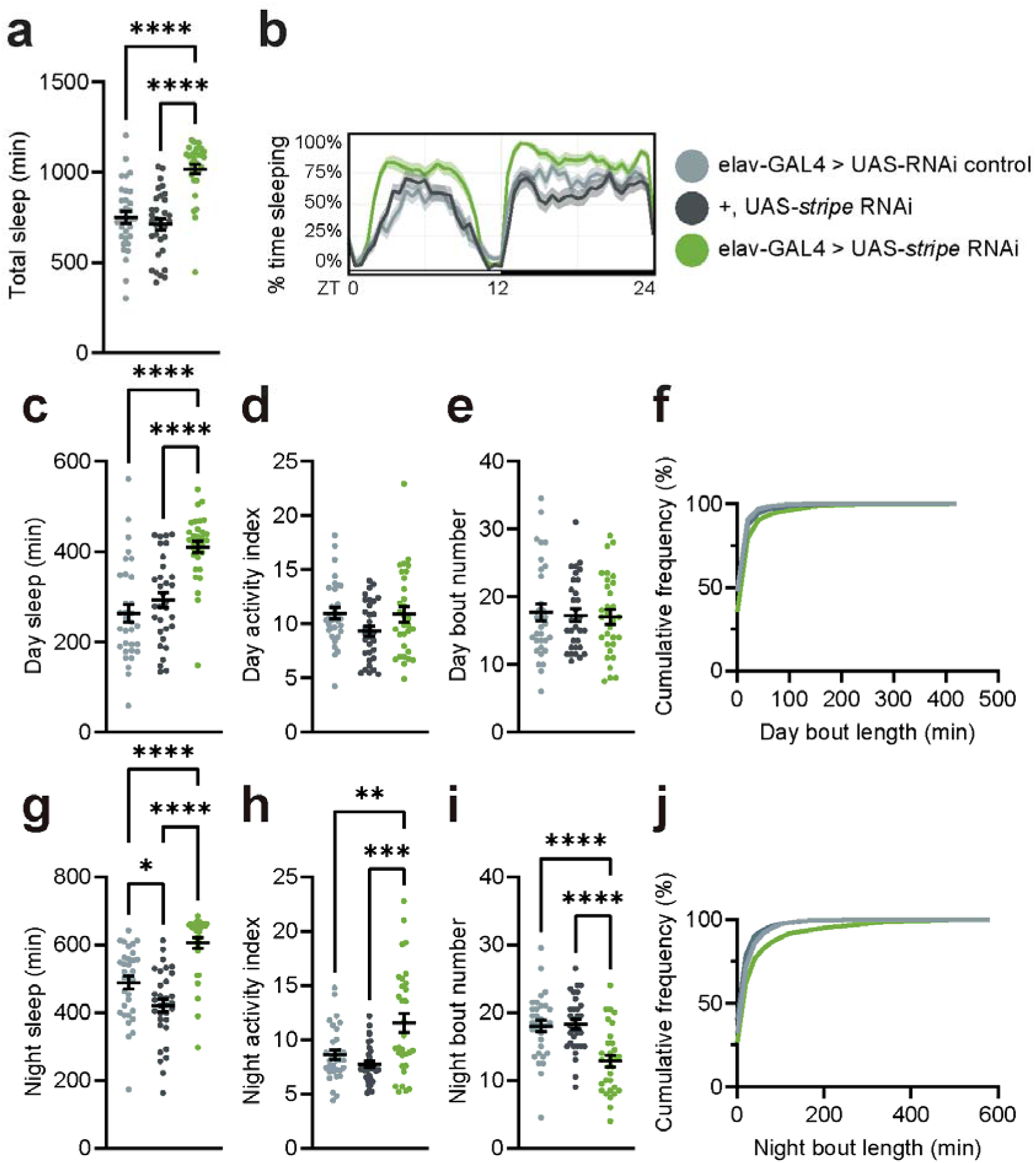
Neuronal *stripe* knockdown increases sleep without reductions in locomotor activity. (a) Total sleep duration across 24 h. (b) Sleep trace, depicting sleep amount (%) in rolling 30 min bins across day (ZT0-12, light) and night (ZT12-24, dark). (c) Daytime sleep duration. (d) Daytime activity index, a measure of locomotor activity (infrared beam breaks per waking minute). (e) Number of daytime sleep bouts. (f) Cumulative frequency of the relative frequency of daytime sleep bouts of increasing duration. (g) Nighttime sleep duration. (h) Nighttime activity index. (i) Number of nighttime sleep bouts. (j) Cumulative frequency of the relative frequency of nighttime sleep bouts of increasing duration. Sleep experiments were run with multibeam DAM monitors. One-way ANOVA with Tukey’s multiple comparisons tests comparing all genotypes. *n,* left to right: 32, 32, 31. ** p ≤ 0.01, *** p ≤ 0.001, **** p ≤ 0.0001.

### Convergent evidence supports EGR2 as an effector gene at the ADO–EGR2 locus

The Cluster 2 variant rs10995311 emerged from our analysis as a candidate hypersomnolence-associated variant. Notably, rs10995311 is in linkage disequilibrium (r² = 0.68 in the EUR population) with rs10995305, a variant previously associated with daytime napping in population GWAS (*9*), providing orthogonal/independent support for the relevance of this locus to sleep-related traits. Chromatin accessibility and promoter capture analyses demonstrated that rs10995311 resides within an open chromatin region that physically contacts the promoter of the distal gene *EGR2* in iPSC-derived neurons (**Figure 1G**). This contact was unambiguously called in iPSC-derived neurons. Candidate contacts in NPCs and astrocytes fell ∼100-200 bp short of the nearest *EGR2*-promoter ATAC-seq peak – within the assay’s resolution limits – and may reflect weaker or technically under-resolved interactions rather than true absence. The neuronal signal is the most robust of the three and is consistent with the sleep phenotype caused by knockdown of *stripe* using a neuron-specific driver.

To further prioritize candidate causal variants at the locus, we annotated rs10995311 and rs10995305 for chromatin state and regulatory features. rs10995311 resides within active promoter-associated chromatin and overlaps numerous predicted transcription factor binding sites, whereas rs10995305 lies predominantly within quiescent chromatin. These observations support rs10995311 as the more likely causal variant underlying the association signal.

Notably, rs10995311 is also a predicted missense variant in exon 1 of *ADO* (Pro39Ala), raising the possibility that *ADO* itself could represent the effector gene at this locus. To address this, we performed *in-silico* analyses of the predicted structural consequences of the amino acid substitution. Multiple prediction algorithms classified the variant as benign or tolerated (AlphaMissense (*18*) = likely benign; CADD phred (*19*) = 6.7 (below the typical ≥15 deleteriousness threshold); PolyPhen-2 (*20*) = benign [0.017]; SIFT (*21*) = tolerated [0.20]; Grantham (*22*) = 27 (conservative)), and examination of five available ADO crystal structures on https://www.rcsb.org/structure (*23*) indicated that Pro39 resides within an unresolved N-terminal disordered region distant from the enzyme active site. Furthermore, HEK293T cells used for structural and inhibitor-binding studies of ADO naturally harbor the P39A variant and produce protein that folds in the canonical cupin conformation and supports hydralazine binding, indicating that the substitution is structurally tolerated (*24*). Finally, neuronal knockdown of *CG7550* (the *Drosophila ADO* ortholog) did not consistently alter sleep (**Supplementary Figure 3**), in contrast to phenotypes observed with *stripe*/*EGR2* knockdown. Thus, while we cannot completely rule out a role for *ADO*, these findings support rs10995311 as a regulatory variant embedded within an *ADO* coding exon that influences *EGR2* expression through neuron-specific chromatin interactions.

### stripe regulates sleep stability and consolidation through olfactory circuits

We next characterized sleep architecture in *stripe* knockdown animals in greater detail. Neuronal *stripe* knockdown increased the proportion of time in long sleep bouts (> 60 min), a feature associated with deep sleep in *Drosophila* (*25*) (**Figure 4A**). Consistent with this, *stripe* knockdown reduced the probability of transitioning from sleep to wake (pWake), indicating increased sleep stability (*26*) (**Figure 4B**). In contrast, measures of wake initiation and maintenance were largely preserved: *stripe* knockdown did not alter sleep latency after lights-on transitions at ZT0 (**Figure 4C**) or the probability of transitioning from wake to sleep (*26*) (pDoze; **Figure 4D**). Sleep homeostasis and circadian rhythmicity also remained intact (**Figure 4E-H**). *stripe* knockdown, therefore, promotes more stable and consolidated sleep without producing broad defects in wakefulness or circadian regulation.

**Figure 4.**
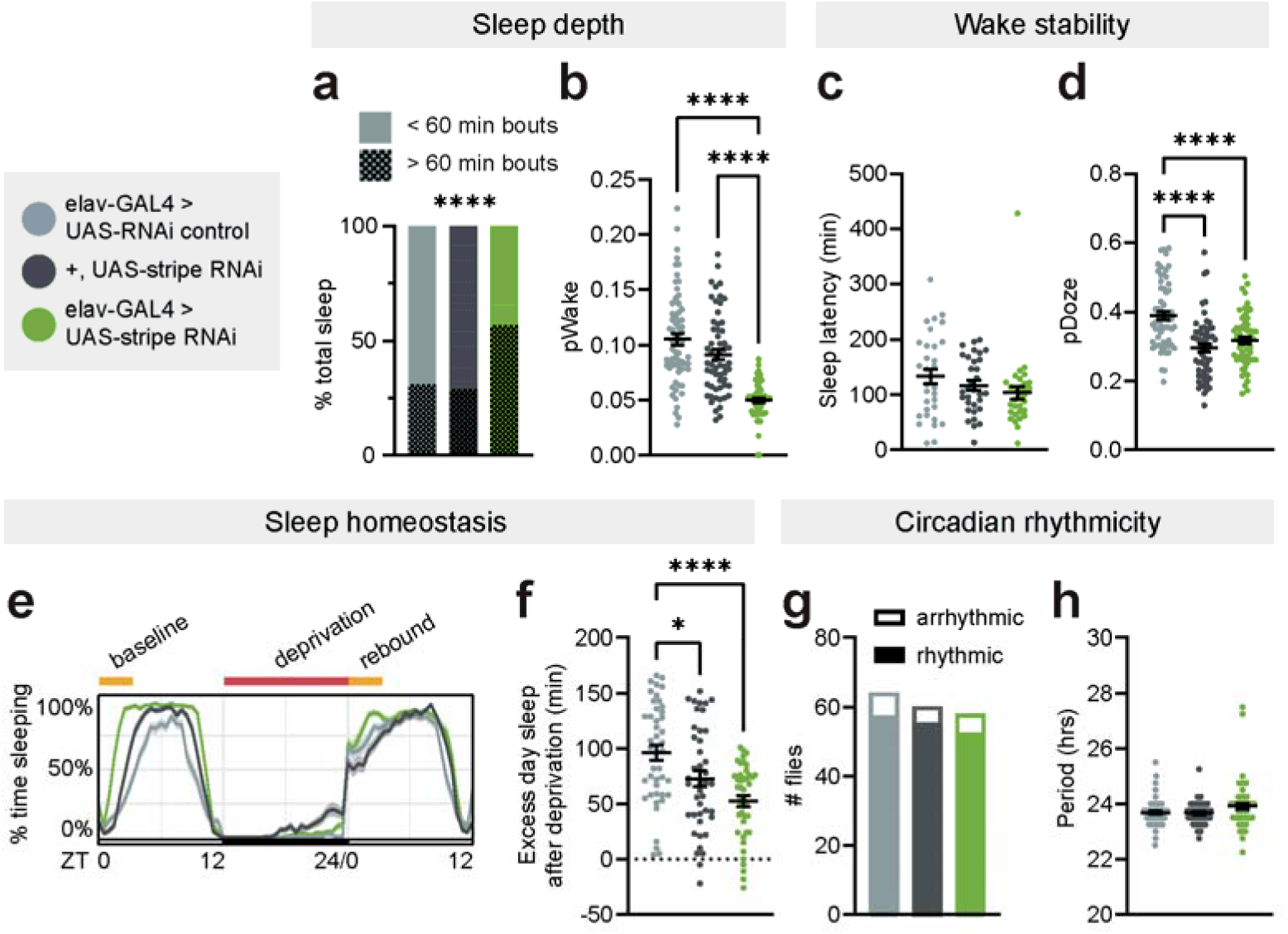
*stripe* knockdown increases sleep depth without affecting waking stability. (a) Percent of total sleep spent in long sleep bouts (> 60 min). (b) Average probability of transitioning from a sleep state to a wake state (pWake) across 24 hrs. (c) Average latency until first sleep bout after daytime light transition at ZT0. (d) Average probability of transitioning from a wake state to a sleep state (pDoze) across 24 hrs. (e) Sleep trace of sleep deprivation experiment. Animals were deprived of sleep overnight (ZT12-24). (f) Excess sleep after sleep deprivation, calculated as recovery sleep – baseline sleep (ZT0-3). (g) Number of animals categorized as rhythmic/arrhythmic under constant conditions. (h) Period length of rhythmic animals under constant conditions. Sleep experiments run with single-beam DAM monitors, except long bout analysis (a) and sleep latency analysis (c) which were run with multibeam DAM monitors. One-way ANOVA with Tukey’s multiple comparisons test comparing all genotypes, except (a) and (g) which were analyzed with Fisher’s exact test. * p ≤ 0.05, **** p ≤ 0.0001.

Both *Drosophila stripe* and human *EGR2* encode transcription factors with diverse roles in neural development and function (*27–31*). Because *stripe* has been implicated in multiple neural populations involved in sensory processing and arousal regulation (*32–36*), we next performed a targeted spatial screen using GAL4 drivers labeling candidate cell populations, including olfactory, mechanosensory, gustatory, circadian, and dopaminergic neurons (**Figure 5A**). Among these drivers, *stripe* knockdown using two broadly expressed olfactory drivers, orco-GAL4 (*37*) and peb-GAL4 (*38*), consistently increased sleep bout duration, whereas most other cell populations produced minimal or no sleep phenotypes (**Figure 5A; Supplementary Figure 4**). Replication using higher-resolution sleep monitoring confirmed that olfactory *stripe* knockdown largely recapitulates the pan-neuronal phenotype (**Figure 5B-K**). Knockdown with either orco-GAL4 or peb-GAL4 increased total sleep and sleep bout duration during both day and night while reducing bout number, consistent with enhanced sleep consolidation. Importantly, locomotor activity during wakefulness was not reduced (**Figure 5F,J**), arguing against nonspecific locomotor impairment. Together, these findings identify olfactory circuits as a major site through which *stripe* regulates sleep duration and consolidation.

**Figure 5.**
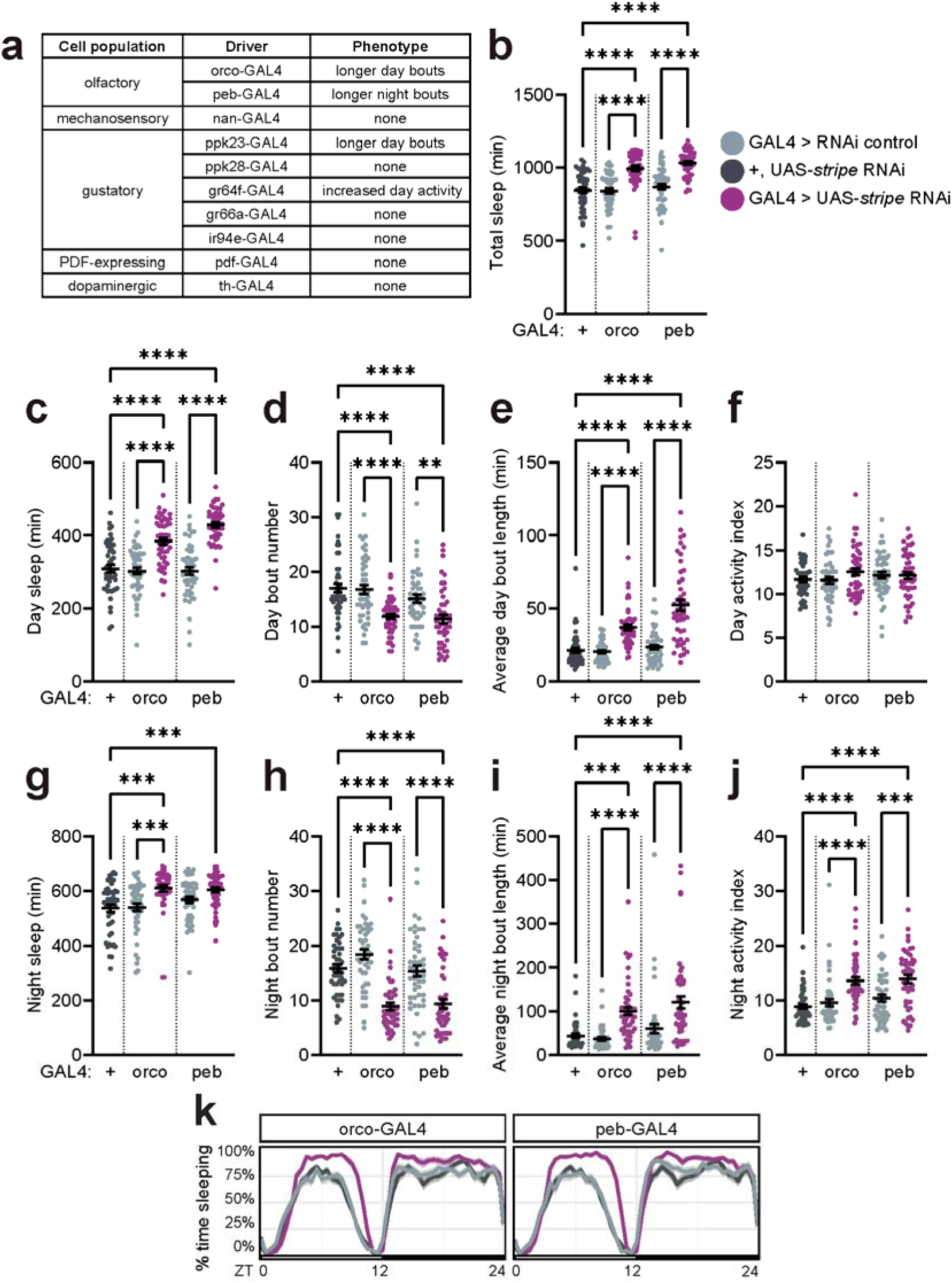
*stripe* is required in olfactory cells for normal adult sleep. (a) Summary of phenotypes found with targeted spatial knockdown screen. Full data can be found in **Supplementary Figure 4**. (b) Total sleep duration across 24 h. (c-f) Daytime sleep measures. (g-j) Nighttime sleep measures. (k) Sleep trace. Sleep experiments were run with multibeam DAM monitors. One-way ANOVA with Šídák’s multiple comparisons test comparing +>UAS-*stripe* RNAi to all genotypes and GAL4>UAS-*stripe* RNAi to their respective GAL4>RNAi control genotypes. *n,* left to right: 48, 47, 48, 48, 48. *** p ≤ 0.001, **** p ≤ 0.0001.

### EGR2 regulates sleep across species

Given the conserved effects of *stripe/EGR2* perturbation on sleep phenotypes in *Drosophila*, we next asked whether *EGR2* similarly regulates sleep in vertebrates by examining the effect of *egr2* mutation in zebrafish. Zebrafish contain two paralogs, *egr2a* and *egr2b*, located on chromosomes 17 and 12, respectively. Using CRISPR-Cas9-mediated mutation, we designed guide RNAs targeting the highly conserved functional domains of both orthologs in zebrafish (*39–41*) (**Supplementary Figure 5**). Consistent with the effects of *stripe* knockdown in *Drosophila*, *egr2ab* mutation increased daytime sleep duration in zebrafish by ∼1.5 hours across the 14-hour daytime period (**Figure 6A,C**). This effect was accompanied by increases in both sleep bout number and length (**Figure 6E,F**). No significant effects were observed for nighttime sleep measures in zebrafish (**Figure 6G-J**). Importantly, waking activity was not altered during either the day or night (**Figure 6B,D,H**).

**Figure 6.**
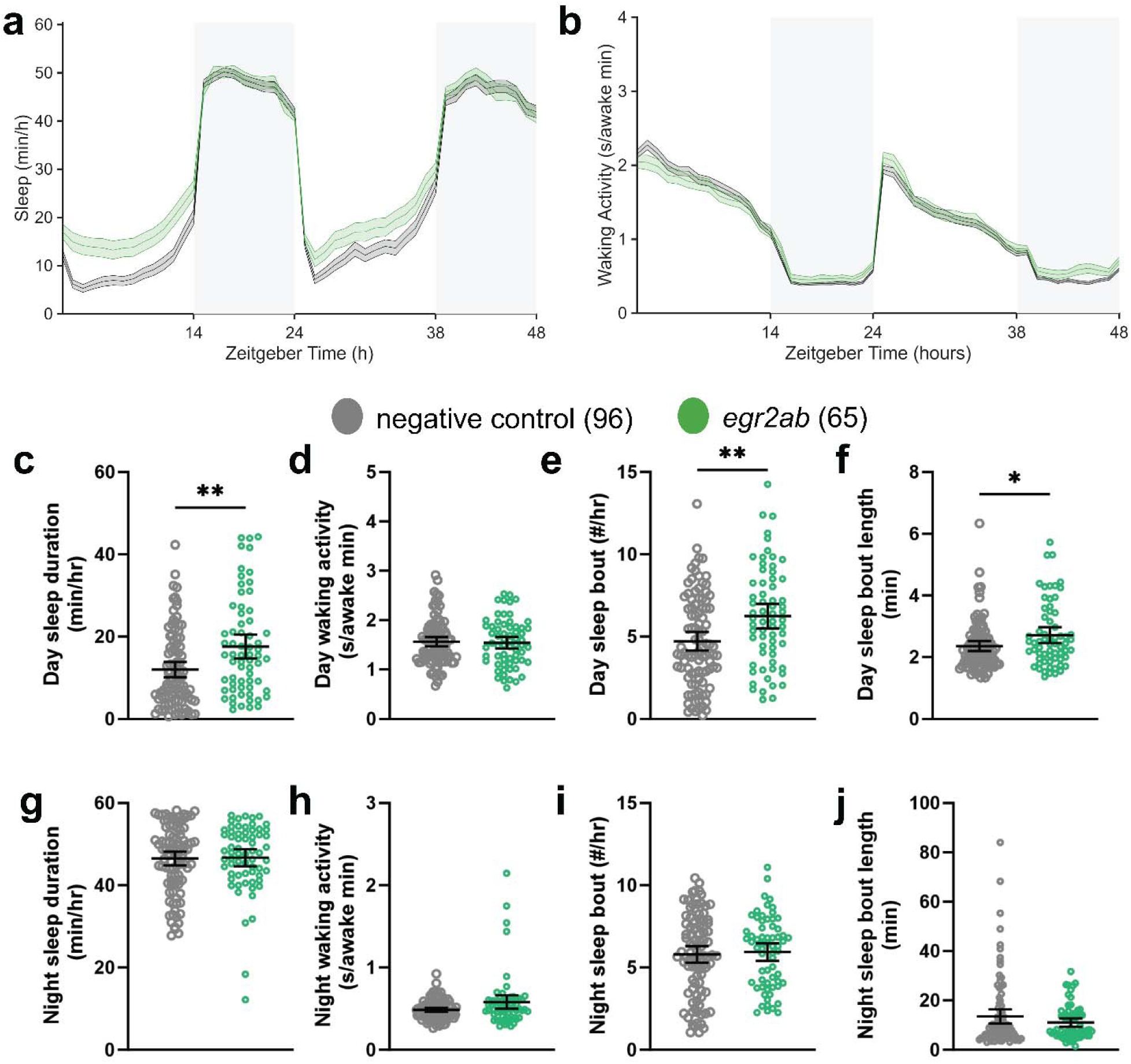
Mutation of *EGR2* orthologs increases daytime sleep in zebrafish. (a) Sleep trace across days 6 and 7 post fertilization in *egr2ab* crispants (n = 65) and negative controls (n = 96). Data are represented as mean ¿ SEM b. Waking activity trace across days 6 and 7 post fertilization. (c-j) Selected sleep and activity parameters in *egr2ab* crispants and negative controls. Data are from two pooled clutches (sibling-matched controls within clutch). * p ≤ 0.05, ** p ≤ 0.01, *** p ≤ 0.001, **** p ≤ 0.0001 determined by non-parametric Mann-Whitney test for all measures except day sleep bout, which was normally distributed and used student’s unpaired t-test.

Because some *egr2ab* crispants displayed abnormalities as they developed (e.g. kinked tail that impacted swimming), only fish that appeared morphologically normal throughout the assay period and showed normal response to touch were included in behavioral analyses (**Supplementary Figure 6A,B; Supplementary Videos 1,2**). A small subset of crispants exhibited intermittent hyperactive bursting behavior and twitching suggestive of seizure-like activity, most prominently at night (**Supplementary Figure 7; Supplementary Videos 1,2**). However, these episodes were not observed in the majority of larvae and did not obscure the overall increase in daytime sleep across the cohort. Together, these findings support a conserved role for EGR2 orthologs in the regulation of sleep across species.

## DISCUSSION

Human genetic studies have demonstrated that sleepiness-related traits, including daytime sleepiness, sleep duration, and daytime napping, are genetically correlated (*7–9*). At the same time, emerging experimental efforts have shown that human sleep-associated loci can be translated into conserved sleep biology through variant-to-gene mapping and cross-species functional validation (*10*, *42*). In the present study, we integrate these complementary approaches by combining multi-trait clustering of sub-threshold sleep GWAS signals (to overcome the relative paucity of genome wide significant signals) with cell type-specific regulatory mapping and experimental validation across species. Using these epigenetically-informed methodologies, we identify *EGR2/stripe* as a conserved regulator of sleep and demonstrate that sleep-regulatory pathways can be recovered from distributed genetic signals that fall below conventional genome-wide significance thresholds.

Sleep traits are highly polygenic, suggesting that relevant pathways may be distributed across many variants of modest effect rather than a small number of loci with large effects. Individual associations below genome-wide significance are difficult to interpret, as any single variant may represent a true but underpowered association or statistical noise. By clustering sub-threshold variants according to their shared relationships with multiple sleep phenotypes, we identified genetic signals more likely to reflect shared biological pathways than isolated associations. Recovery of *CADM2* within Cluster 2 – previously identified as a hypersomnolence effector through TAD-wise V2G of genome-wide significant loci (*10*) – provides an internal methodological control confirming that multi-trait clustering of sub-threshold variants captures *bona fide* hypersomnia biology beyond the genome-wide-significance threshold. Thus, rather than treating variants that fail to achieve genome-wide significance as background noise, our findings suggest that latent structure within polygenic sleep-trait architecture can reveal pathways underlying sleep and sleepiness.

Among the strongest candidates emerging from this analysis was *EGR2/stripe*. Multiple independent RNAi lines targeting *stripe* increased sleep in *Drosophila*, and mutation of zebrafish *egr2* orthologs similarly increased sleep. The convergence of human genetics, chromatin architecture, *Drosophila* functional screening, and vertebrate validation provides strong evidence that *EGR2* is involved in evolutionarily conserved pathways regulating sleep. Importantly, the sleep phenotype produced by *stripe* knockdown was characterized not only by increased sleep duration but also by increased sleep consolidation and reduced probability of transitioning from sleep to wakefulness, while leaving locomotor activity, circadian rhythmicity, and sleep homeostasis largely intact. Together, these findings argue against a nonspecific effect on behavioral state and instead support a more specific role for *EGR2/stripe* in regulating sleep stability and propensity.

*EGR2* has not been previously implicated in hypersomnolence-related traits, but evidence supports a broader role in sleep-wake regulation. *EGR2* is among a small number of transcription factors consistently induced by sleep deprivation across multiple brain regions and species, like established activity-dependent sleep genes such as *Homer1a* (*28*, *43*). Our work indicates *EGR2* is not simply an output of sleep deprivation but is an active participant in sleep-wake stability. More generally, *EGR2* functions as an immediate-early transcription factor involved in activity-dependent transcriptional programs, neuronal plasticity, and circuit remodeling (*30*). Sleep is tightly coupled to neural plasticity, raising the possibility that *EGR2* participates in transcriptional programs linking waking neural activity to subsequent sleep need. Future studies defining the cellular targets and transcriptional networks regulated by *EGR2* during sleep and wakefulness may provide insight into the mechanisms through which this pathway influences sleep behavior.

Our findings further support *EGR2* as a likely effector gene at the *ADO–EGR2* locus. The candidate variant rs10995311 resides within an open chromatin region that physically contacts the *EGR2* promoter in neurons and was prioritized through cell-type-specific V2G mapping, while structural analyses suggest that the predicted *ADO* missense variant is unlikely to alter protein function. The observation that this chromatin interaction was identified in neuronal datasets, along with the sleep phenotypes produced by perturbation of *EGR2* orthologs in flies and fish and the absence of hypersomnolence with knockdown of the *ADO* ortholog in flies, collectively provide evidence supporting *EGR2* as a significant mediator of this association signal. These results are consistent with emerging evidence that coding exons can function as regulatory elements controlling distal genes (*44*). raising the possibility that rs10995311 represents a regulatory variant embedded within an *ADO* coding exon that influences *EGR2* expression through long-range chromatin interactions.

An unexpected finding from our study was that knockdown of *stripe* in olfactory neurons was sufficient to recapitulate the sleep phenotype observed following pan-neuronal knockdown. Sensory processing plays an important role in sleep regulation, as sleeping animals must continuously balance responsiveness to salient environmental stimuli against the maintenance of stable sleep states (*45*, *46*). Prior work has implicated olfactory circuits in sleep regulation across species (*47*, *48*). supporting a connection between sensory processing and sleep control. The identification of olfactory neurons as a major site of *stripe* action suggests that sensory circuits may represent an important node through which *EGR2/stripe* influences sleep behavior. Notably, patients with narcolepsy have been found to exhibit reduced ability to detect and identify odors, further suggesting a link between olfaction and hypersomnolence (*49–51*). Whether *stripe* regulates olfactory circuit development, neuronal excitability, sensory responsiveness, or other aspects of sensory function remains unknown. Regardless, these findings support the idea that altered sensory processing may represent an underappreciated contributor to increased sleep propensity and excessive sleepiness.

Several limitations should be considered when interpreting these findings. First, the clustering approach relies on sub-threshold genetic associations, which inevitably contain a mixture of true signals and statistical noise. Although conservative cluster assignment thresholds were used, some candidate variants are likely to represent false positives. Second, the identified variant cluster likely reflects broader hypersomnolence-and sleep propensity-related biology rather than idiopathic hypersomnia specifically, reflecting both phenotypic overlap among sleep traits and the limitations of current population-scale sleep phenotyping approaches. Third, variant-to-gene mapping remains limited by the availability and resolution of chromatin accessibility and interaction datasets, and the causal genes and regulatory mechanisms underlying many associated loci remain uncertain. Finally, the *Drosophila* screen relied on RNAi-mediated knockdown, and although multiple independent RNAi lines were used for key candidates, off-target effects cannot be completely excluded from the broader screen.

Excessive sleepiness is a highly polygenic trait, suggesting that much of its underlying biology resides within distributed genetic signals that are difficult to interpret individually. Our findings demonstrate that integrating multi-trait genetic architecture with functional genomics and cross-species validation can reveal sleep-regulatory pathways hidden within this polygenic landscape. More broadly, these results suggest that latent structure across related sleep phenotypes may provide a powerful framework for uncovering mechanisms that contribute to hypersomnolence-related traits.

## METHODS

### GWAS data sources and variant selection

Sub-threshold variants (P ≤ 5 × 10^-^□) were extracted from three large-scale sleep GWAS: daytime napping(*9*) (UK Biobank, N = 452,633), daytime sleepiness(*7*) (UK Biobank, N = 452,071), and sleep duration(*8*) (UK Biobank, N = 446,118). Five primary traits were used as index phenotypes for variant extraction. All variants were mapped to hg38/GRCh38 coordinates in the format CHR_POS_REF_ALT. Indels were excluded; only biallelic SNPs were retained. (**Supplementary Table 1**)

### Variant filtering and LD pruning

Variant filtering was performed using custom R scripts (choose_variants.R). Variants were first clumped by genomic proximity using a 500 kb window (snp_clump() function). LD pruning was then performed using LDlinkR SNPclip with the following parameters: r² threshold = 0.05, MAF threshold = 0.001, reference population = EUR (1000 Genomes Phase 3). After pruning, trait availability was assessed across the 18 UK Biobank sleep phenotypes using count_traits_per_variant(). Variants with >60% missing traits were flagged for proxy replacement. LD proxies were identified using TopLD API (primary) with LDlinkR fallback (choose_proxies() function), selecting the proxy with maximum trait coverage. Variant IDs were mapped to rsIDs using dbSNP build 155 VCF (generate_varid_to_rsid_map_file.R). The final input matrix contained 4,254 independent variants (**Supplementary Table 2**).

### Sleep trait z-score extraction

For each retained variant, association z-scores (BETA/SE) were extracted from summary statistics of 18 UK Biobank sleep phenotypes:

- **Self-reported (9 traits):** sleep duration, daytime napping, daytime sleepiness, chronotype, ease of getting up, long sleep, short sleep, snoring, insomnia.
- **Accelerometer-derived (8 traits):** sleep duration (ACC), sleep efficiency, number of sleep episodes, L5 timing, M10 timing, diurnal inactivity, sleep duration variability, sleep midpoint.
- **Clinical (1 trait):** obstructive sleep apnea (OSA).

Z-scores were scaled by per-trait median sample size: z_scaled = z / √(medN_trait) × mean(√(medN_all_traits)). Missing z-scores were imputed using trait median values or LD proxy z-scores where available. The z-score matrix was expanded into a non-negative format by separating positive and negative components (N variants × 2M trait columns), following.(*11*)

### Bayesian non-negative matrix factorization (bNMF)

The non-negative z-score matrix was decomposed using the BayesNMF.L2EU algorithm(*11*) following with automatic relevance determination (ARD) and half-normal priors on the W (variant × cluster) and H (cluster × trait) matrices. The algorithm was initialized with K=20 and run 1,000 times with convergence tolerance 1 × 10^-^□ and maximum 1,000 iterations per run. The optimal number of clusters was determined by comparing negative log-evidence across K=3-7 over the 1,000 runs (run_bNMF.R). K=4 was selected over the statistically optimal K=5 because the K=5 hypersomnia signal was distributed across multiple clusters and 80.6% of K=5 hypersomnia-cluster V2G genes were recovered in the K=4 Cluster 2 set, with K=4 nominating 68 additional candidates (**Supplementary Figure 1**) (see Results). Variants were assigned to clusters using a 95th percentile weight threshold on the W matrix.

### Variant-to-gene (V2G) mapping

#### cRE definition

Cis-regulatory elements (cREs) were defined per cell type as the intersection of ATAC-seq peaks (open chromatin regions, OCRs) with 3D chromatin architecture features, and with promoters (-1,500/+500bp of TSS) defined by GENCODE v47.

#### Cell types

V2G was performed using in-house chromatin data from three neural cell types:

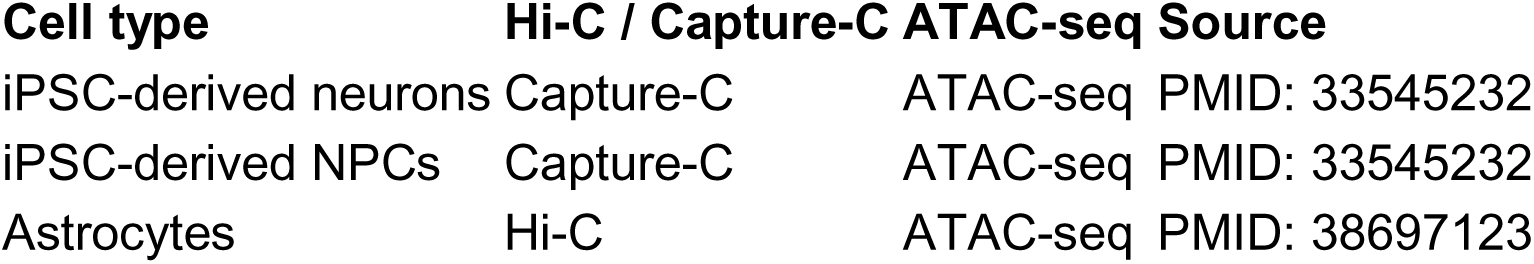

### ADO-EGR2 locus characterization

#### Functional annotation

Variant-level annotation for rs10995311 and rs10995305 was performed using myvariant.info API to retrieve CADD phred scores, PolyPhen-2, SIFT, AlphaMissense predictions, ENCODE regulatory marks (H3K4me3, H3K27ac, DNase), ChromHMM chromatin state frequencies, and transcription factor binding site counts. Grantham scores were used to assess amino acid substitution severity.

#### Protein structure analysis

ADO protein structure was assessed using five experimental crystal structures in the Protein Data Bank (PDB: 7REI, 7LVZ, 8UAN, 9DXV, 9DMA) spanning human and mouse ADO in apo and inhibitor-bound states. The position of Pro39 (rs10995311) was evaluated relative to the cupin superfamily beta-barrel fold and the 3-His facial triad active site (His112, His114, His193). AlphaFold predicted structure (UniProt Q96SZ5) was used for regions not resolved in crystal structures.

#### eQTL analysis

eQTL lookup was performed in GTEx v8 (all 49 tissues), eQTLGen blood meta-analysis (N = 31,684), MetaBrain cortex meta-analysis, and SingleBrain single-cell brain atlas. Both direct variant queries and LD proxy queries (r² > 0.6, EUR) were used to maximize coverage.

### Drosophila RNAi screen

#### Drosophila husbandry

Unless otherwise specified, flies were raised and maintained on standard molasses food (8.0% molasses, 0.55% sugar, 0.2% Tegosept, and 0.5% propionic acid) at 25°C on a 12h:12h light:dark (LD) cycle. 5-7 d old female flies were used in all experiments.

#### Fly strains

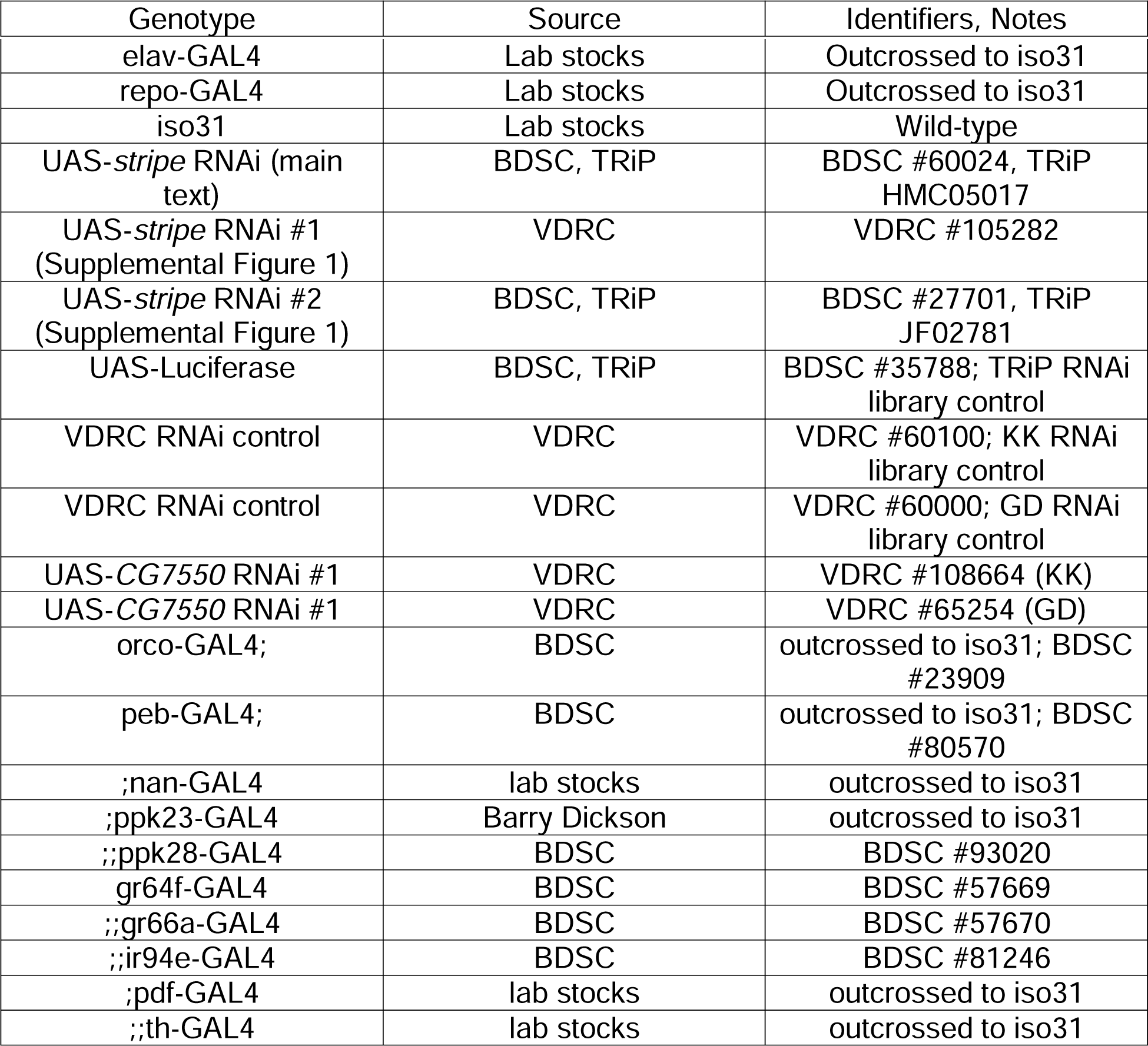

#### Sleep phenotyping

##### Sleep assays

Flies were anesthetized on CO_2_ pads (Genesee Scientific, #59-114) and loaded into individual glass tubes supplied with 5% sucrose, 2% agar food. Tubes were loaded into Drosophila Activity Monitoring Systems (Trikinetics) and flies were monitored for at least 2 d. Activity was measured in 1 min bins and sleep was defined as 5 min of inactivity, except for long bout analyses for which sleep was defined as 60 min of inactivity. Activity indices were measured as number of beam breaks per 1 min of waking activity. Data were analyzed using Rethomics (*52*), except for pWake/pDoze analyses which were analyzed using SCAMP (*53*)/

##### Sleep latency

Sleep latency was measured as time lapsed until animals’ first sleep bout following lights on at ZT0. Animals that were not awoken by the light transition were excluded.

##### Sleep homeostasis

Following baseline day sleep measurement from ZT0-12, flies received a 2 s stimulation randomly within every 20 s window from ZT12-24 using a mechanical vortexer (Fisher Multitube Vortexer, Catalog # 02-215-450).

##### Circadian rhythmicity

Flies were collected 5-7 d after eclosion and entrained to a 12 h:12 h LD cycle for 4 d. They were then loaded into the DAM system and entrained to a 12h:12h LD cycle for 1 d before transferred to constant darkness (DD). Locomotor activity was measured for 5 d in DD. Locomotor rhythmicity was analyzed using SCAMP (*53*). Animals with a rhythmicity index of at least 0.195 were defined as rhythmic, and period length was calculated only from animals categorized as rhythmic.

##### PCA and clustering

Using GraphPad Prism, PCA was run on 9 sleep measures for each genotype (average day sleep duration, night sleep duration, total sleep duration, day bout number, night bout number, day bout length, night bout length, day activity index, night activity index). Measures were standardized prior to analysis such that data were scaled to have a mean of 0 and a standard deviation of 1. Principal components (PCs) were selected such that they explained at least 65% of the variance of the dataset. Following PCA, k-means clustering was applied to PC1/PC2 values to identify groups of genotypes with shared characteristics. 123 was used as a random seed. The number of clusters was chosen by visual inspection of the scree plot and biological coherence of the heatmap of sleep measures.

### Zebrafish validation

#### CRISPR mutation of *egr2a* and *egr2b*

egr2ab crispants were generated following previously published protocols (*42*, *54*). Highly specific guide RNAs (gRNAs) and target-specific primers were designed using the online tool CRISPOR (http://crispor.tefor.net/) with the reference genome set to “NCBI GRCz11” and the protospacer adjacent motif (PAM) set to “20bp-NGG-Sp Cas9, SpCas9-HF, eSpCas9 1.1.” gRNAs with stringent specificity score >95% with 0 predicted off-targets for sequences with up to 3 mismatches were selected. gRNAs with Lindel (in/del prediction) scores >75 were prioritized (*41*, *55*). gRNAs were designed to target all transcripts identified by ensemble and a conserved protein domain critical to function to increase the likelihood of generating a non-functional protein (*41*). Pre-formed ribonuclear protein (RNP) complexes containing the gRNA:tracr and spCas9 HiFi enzyme (IDT) were injected at the single-cell stage as previously described (*54*). Injections of scrambled (non-targeting) gRNA and gene-specific gRNA were performed in embryos from the same breeding event (clutch). The following Alt-RT^TM^ gRNAs were ordered from IDT (https://www.idtdna.com/) [guide+ *PAM*]: egr2a CTCCGGTCGCATCCCTCGGC *CGG*; egr2b: TCCATCGACTCCCAGTACCC *GGG*.

#### DNA extraction and PCR for genotype validation

DNA extraction was performed per the manufacturer’s protocol (Quanta bio, Beverly, MA) immediately following completion of the sleep assay, as described previously (*39*, *42*). gRNAs were designed to disrupt a restriction enzyme site such that effective mutation could be detected by a lack of restriction enzyme cutting. For standard primers, mutation rates were calculated as the ratio of the uncut:cut amplicon. Select samples were Sanger sequenced to confirm the on-target mutation. Primers used for genotype validation are as follows:

*egr2a* Forward: 5’-CACTGAAACGGCTTGTGTCC-3’;

*egr2a* Reverse: 5’-GCTGCCAAAGGGACAACATG-3’;

*egr2b* Forward: 5’-GAGGTGGAAAGGAACGCAGA-3’;

*egr2b* Reverse: 5’-TTCCCTCTTCCAGATGGCCT-3’

EagI restriction enzyme was used for egr2a and BslI was used for egr2b genotype validation.

#### Data collection and analysis for sleep phenotyping

Sleep and activity tracking were captured using automated video tracking as previously described (*56*, *57*). CRISPR mutants and scramble-injected sibling controls were assessed for gross morphological deficits, and healthy larvae were pipetted into individual wells of a 96-well plate and placed into a Zebrabox (ViewPoint Life Sciences) for automated video monitoring. Genotypes were placed into alternating rows to minimize location bias within the plate. All animals were allowed to acclimate to the Zebrabox for approximately 18 hours before beginning continuous data collection for 48 hours starting at lights on (9am) on day 6 post fertilization. At the conclusion of each assay, all fish were assessed for morphological deficits and tapped with a P10 pipette tip to ensure normal response. Abnormal larvae were removed from analyses. Two biological replicates were run using different clutches (sibling-matched within clutch) of embryos and well placement was flipped for each experiment to minimize location bias across experiments. MATLAB scripts and full protocol for sleep/activity analysis can be found at DOI: <u>10.21769/BioProtoc.4313</u>.

### Statistical analysis

All statistical analyses were performed using R or GraphPad Prism. Details on sample sizes and statistical tests are denoted in figure legends. Dot plots represent mean +/-SEM unless otherwise specified. For all figures, * p ≤ 0.05, ** p ≤ 0.01, *** p ≤ 0.001, and **** p ≤ 0.0001.

## Funding

NIH T32GM008076 (KM)

NIH T32HL007953 (KM)

NIGMS K12GM081259 (EL)

NIH T32HL170968 (AZ)

NIH R35HG011959 (AC)

NIH R01HL143790 (SFG) NIH

R01HL178074 (SFG)

Daniel B. Burke Endowed Chair for Diabetes Research (SFAG)

NIH R01NS135075 (MSK, AC)

NIH R35NS137329 (MSK)

Linda Pechenik Montague Award (MSK)

Burroughs Wellcome Career Award for Medical Scientists (MSK)

## Author contributions

Conceptualization: KM, KT, AZ, AC, SFAG, MSK

Investigation: All authors

Writing – Original Draft: KT, KM, AC, AZ, SFAG, MSK

Writing – Review and Editing: All authors

Project Supervision and Funding: MSK, SFAG

## Data and code availability

All bNMF analysis scripts are available at https://github.com/gwas-partitioning/bnmf-clustering. Raw GWAS summary statistics were accessed from https://sleep.hugeamp.org/downloads.html and https://ftp.ebi.ac.uk/pub/databases/gwas/. V2G chromatin datasets are deposited at E-MTAB-9159, E-MTAB-9087, GSE241592, GSE241591.

## SUPPLEMENTARY FIGURES

**Supplementary Figure 1.**
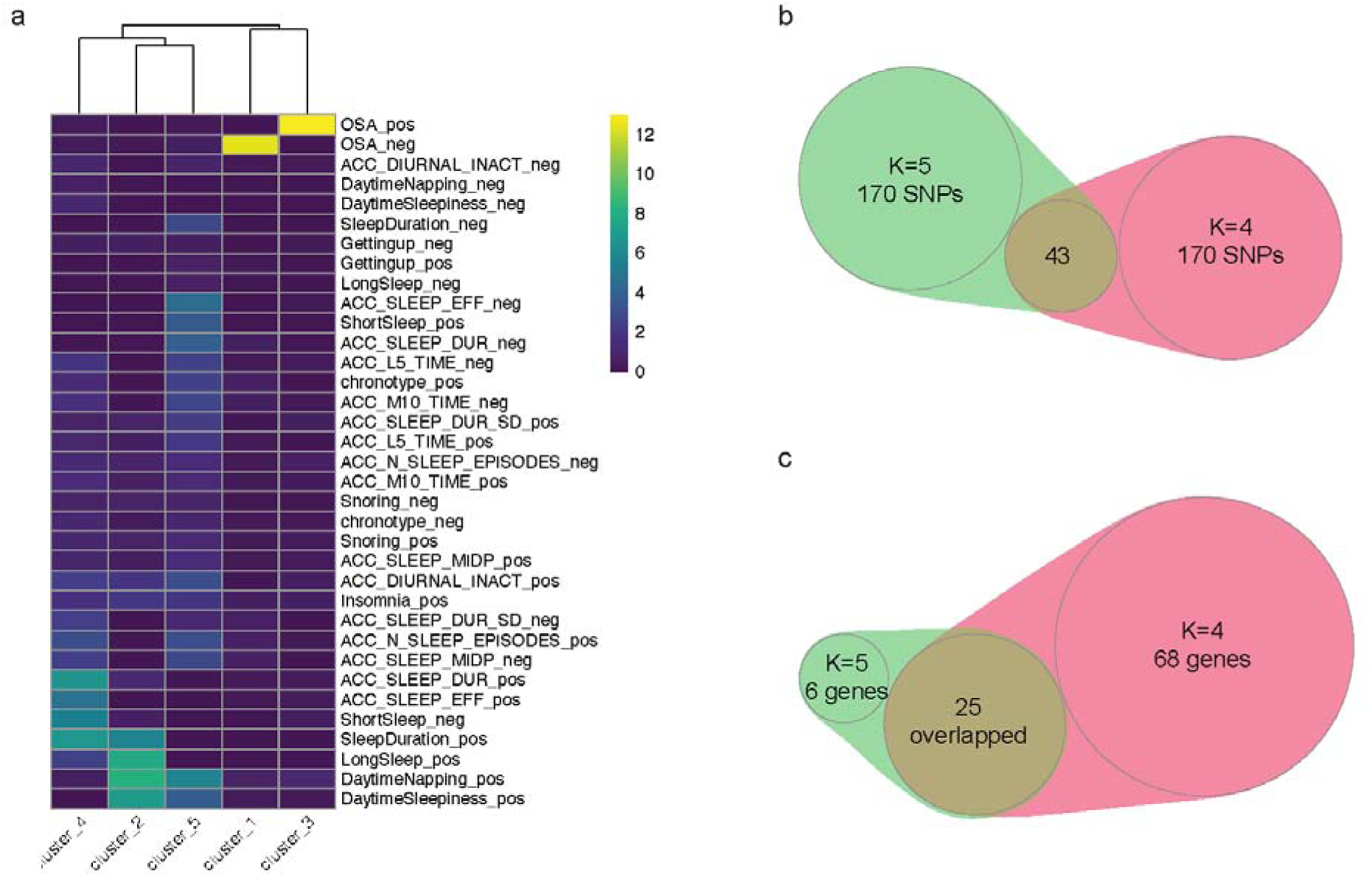
K=5 bNMF solution comparison. (a) Trait weight heatmap (H matrix) for K=5: 5 clusters × 18 traits, z-scores displayed. (b) number of Cluster 2 SNPs under K=4 vs. K=5. (c) V2G gene nomination comparison: number of Cluster 2 genes nominated under K=4 vs. K=5.

**Supplementary Figure 2.**
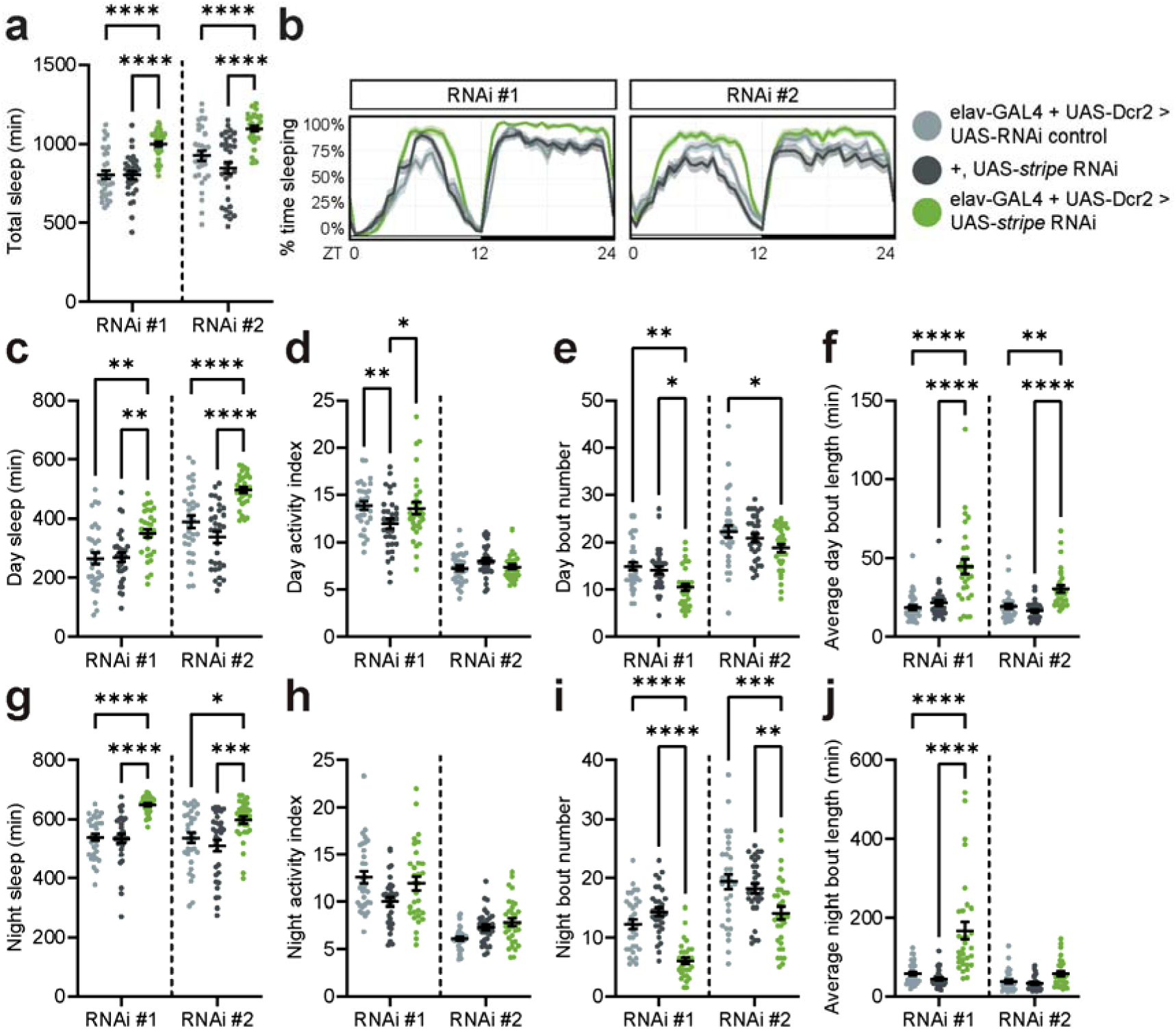
Multiple RNAi against *stripe* produce long-sleeping phenotypes when expressed in neurons. Knockdown of *stripe* using RNAi #1 (VDRC #105282) or RNAi #2 (TRiP #27701). (a) Average total sleep across 24 hrs. (b) Sleep traces, depicting sleep amount (%) in rolling 30 min bins across day (ZT0-12, light) and night (ZT12-24, dark). (c-f) Daytime sleep measures. (g-j) Nighttime sleep measures. * p ≤ 0.05, ** p ≤ 0.01, *** p ≤ 0.001, **** p ≤ 0.0001.

**Supplementary Figure 3.**
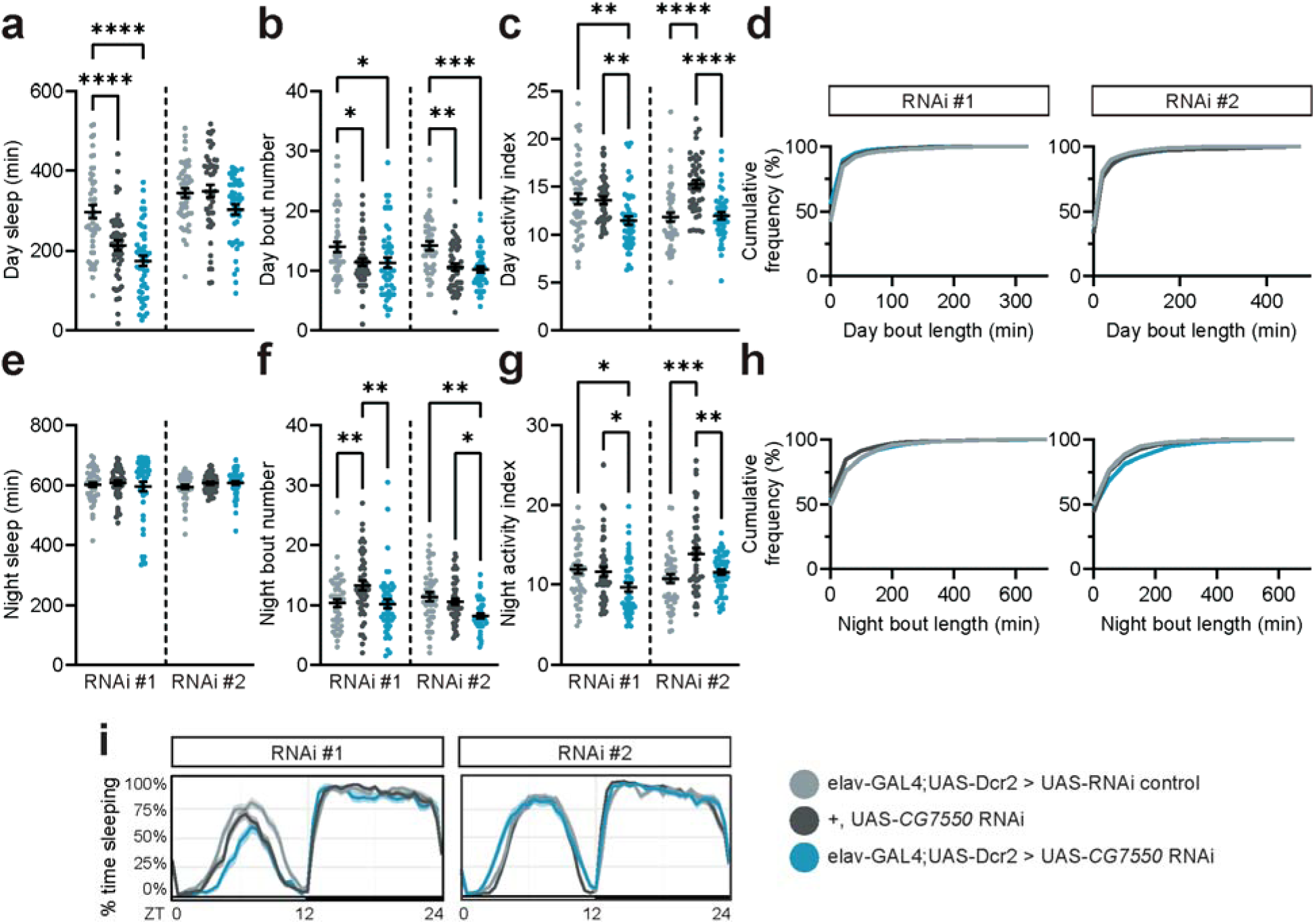
Knockdown of *Drosophila ADO* homolog, *CG7550*, does not increase sleep. (a-d) Daytime sleep measures. (e-h) Nighttime sleep measures. (i) Sleep traces, depicting sleep amount (%) in rolling 30 min bins across day (ZT0-12, light) and night (ZT12-24, dark). Sleep experiments were run with multibeam DAM monitors. One-way ANOVA with Tukey’s multiple comparisons tests comparing all genotypes. *n,* left to right: 32, 32, 31. * p ≤ 0.05, ** p ≤ 0.01, *** p ≤ 0.001, **** p ≤ 0.0001.

**Supplementary Figure 4.**
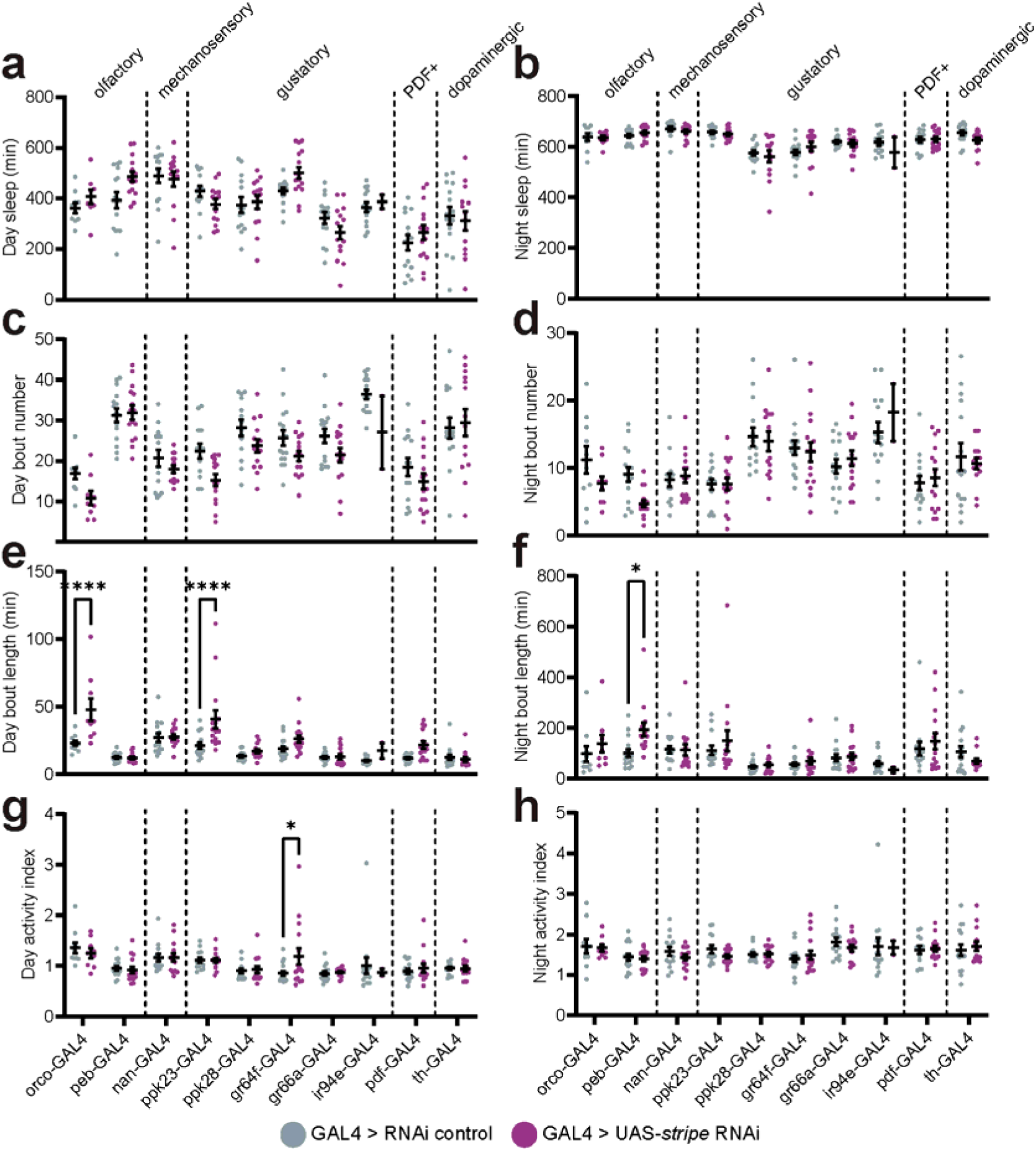
Full targeted spatial knockdown screen data. (a, b) Day and night sleep duration. (c, d) Day and night bout number. (e, f) Day and night average bout length. (g, h) Day and night activity index. Sleep experiments were run with single beam DAM monitors. Two-way ANOVA with Šídák’s multiple comparisons test comparing GAL4>UAS-*stripe* RNAi to their respective GAL4>RNAi control genotypes. *n,* left to right: 10, 9, 15, 15, 14, 14, 13, 15, 15, 15, 16, 16, 15, 15, 14, 2, 14, 15, 15, 14. *p ≤ 0.05, **** p ≤ 0.0001.

**Supplementary Figure 5.**
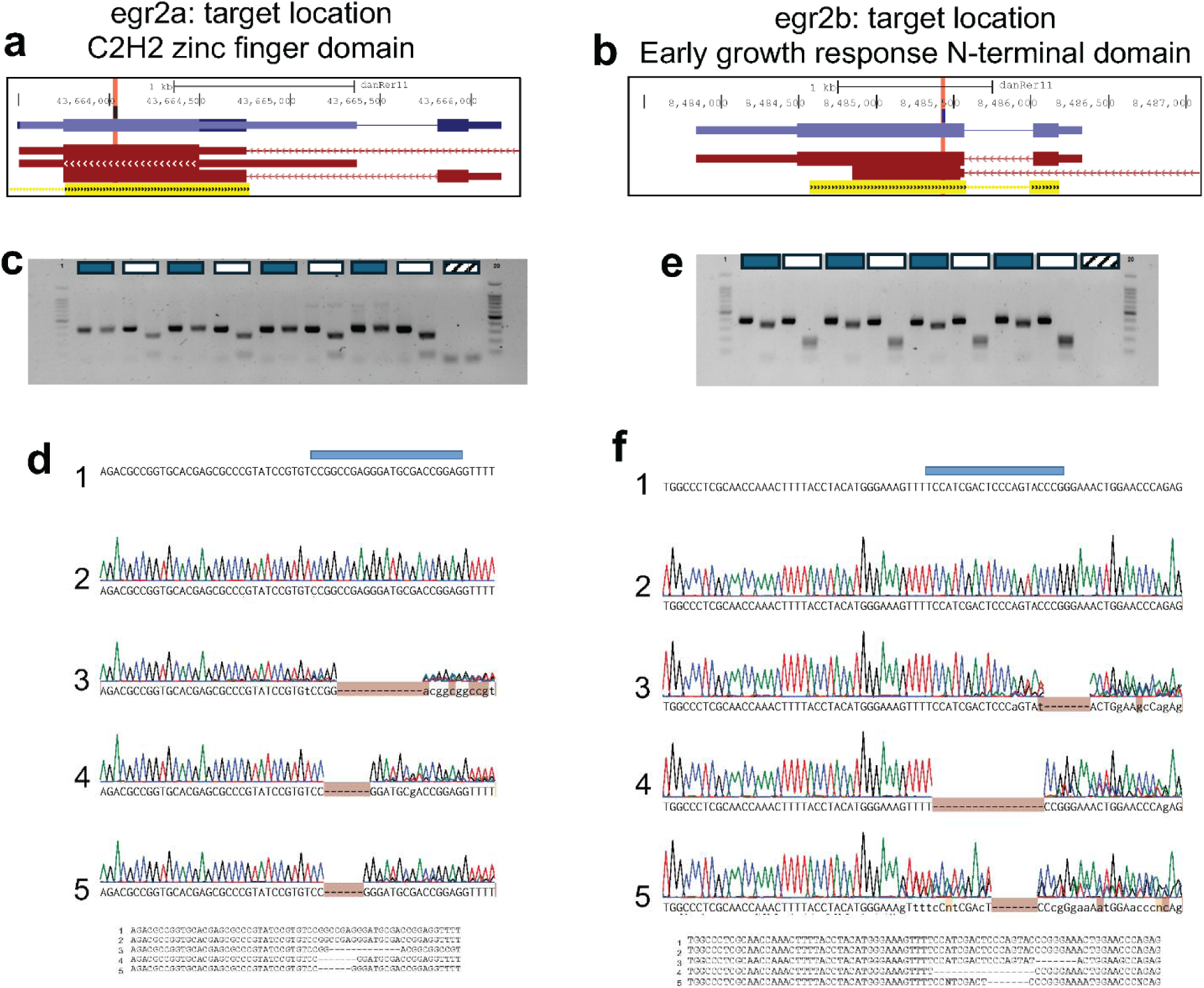
CRISPR mutation validation for *egr2ab* crispants. (a) gRNA location (red line) targets the highly conserved C2H2 zinc finger domain of *egr2a*. (b) gRNA targets the highly conserved Early growth response N-terminal domain of *egr2b*. (c) PCR validation of the on-target mutation for the *egr2a* guide. (d) Representative Sanger traces showing consequence of the mutation of *egr2a* in 2 negative controls (1-2) and 3 crispants (3-5). (e) PCR validation of the on-target mutation for the *egr2b* guide (showing same fish as in (c) given these are double *egr2ab* crispants). (f) Representative Sanger traces showing consequence of the mutation of *egr2b* in 2 negative controls (1-2) and 3 crispants (3-5). In (c) and (e), blue bars represent crispants and white bars represent negative controls. Hashed bars represent primer-only water controls. In (e) and (f), the blue bar on top indicates the gRNA. All crispants sequenced had severe phenotypes indicating each mutation was deleterious.

**Supplementary Figure 6.**
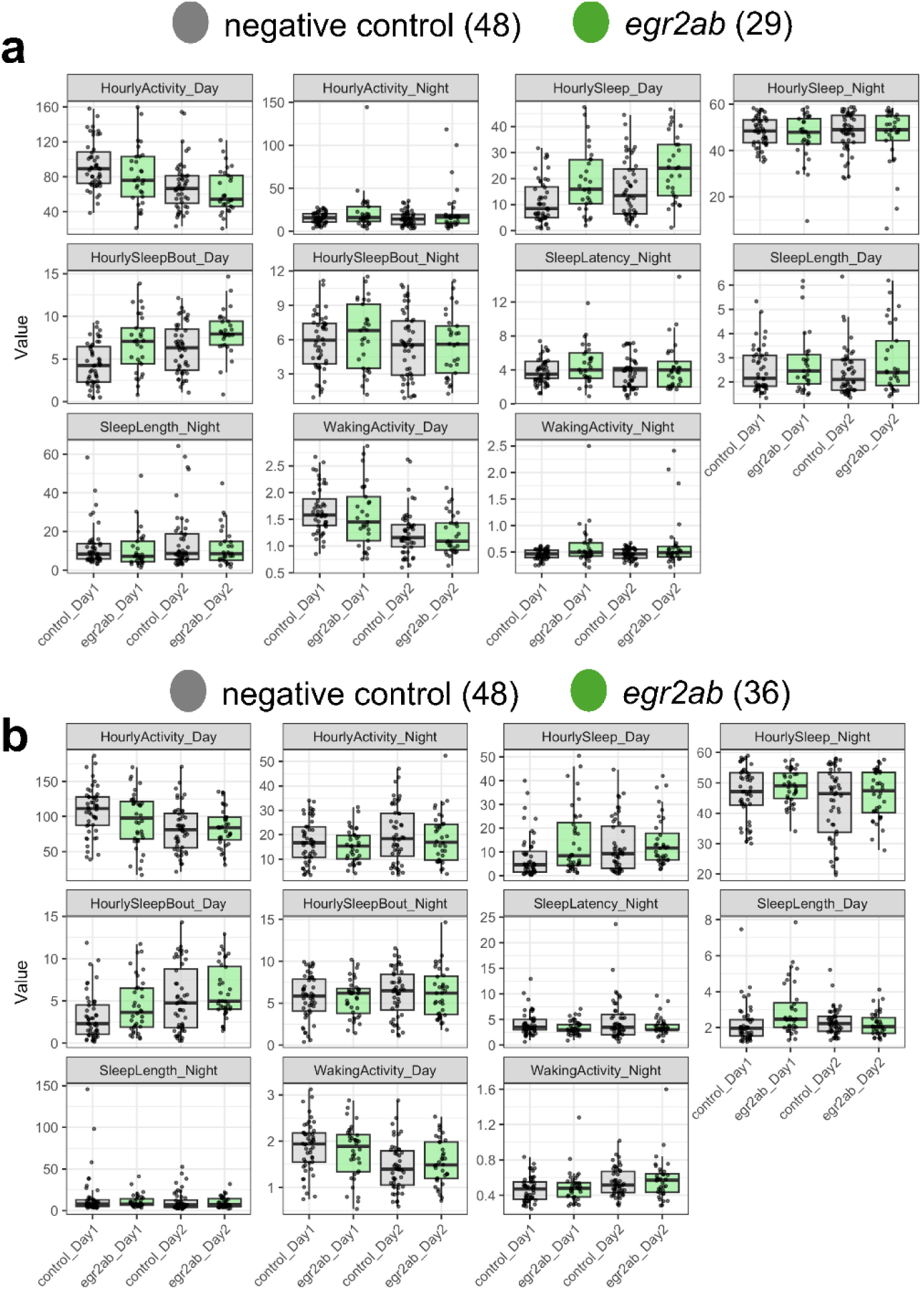
All behavioral measures of *egr2ab* crispants. All behavioral measures from clutch 1(a) and clutch 2 (b). Data are presented as box plots (median and quartiles) along with individual fish for both days measured (6 and 7 dpf). Sample size is indicated next to genotype.

**Supplementary Figure 7.**
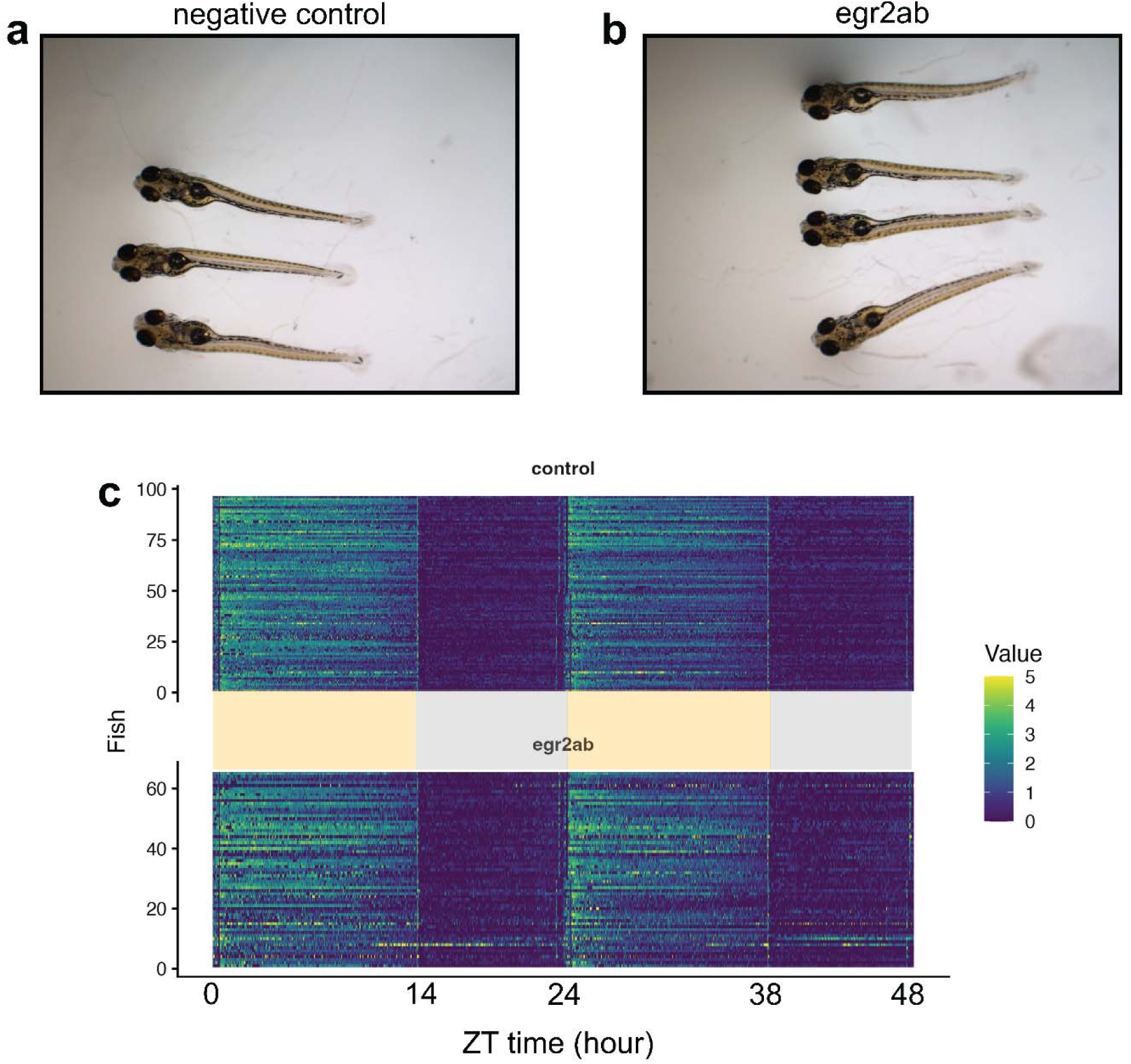
Examples of larvae used in sleep assay. Images showing 3 negative controls (a) and 4 *egr2ab* crispants (b) following the conclusion of the sleep assay at 8dpf. Swim bladders are inflated and normal morphological characteristics are observed. (c) Activity (seconds/min) for all measured fish (n = 96 negative controls and 65 *egr2ab* crispants). Each y-axis bin is 1 minute. Hyperactive bouts (yellow intensity values) can be observed at night (gray boxes) for a few *egr2ab* larvae. Yellow shaded boxes represent day (light-on).

## TABLES

**Table 1**. Full V2G results: all Cluster 2 variants, LD proxies, nominated genes, cell type support.

